# FGF21 protects against hepatic lipotoxicity and macrophage activation to attenuate fibrogenesis in nonalcoholic steatohepatitis

**DOI:** 10.1101/2022.09.20.508654

**Authors:** Cong Liu, Milena Schönke, Borah Spoorenberg, Joost M. Lambooij, Hendrik J.P. van der Zande, Enchen Zhou, Maarten E. Tushuizen, Anne-Christine Andréasson, Andrew Park, Stephanie Oldham, Martin Uhrbom, Ingela Ahlstedt, Yasuhiro Ikeda, Kristina Wallenius, Xiao-Rong Peng, Bruno Guigas, Mariëtte R. Boon, Yanan Wang, Patrick C.N. Rensen

## Abstract

Analogues of the hepatokine FGF21 are in clinical development for type 2 diabetes and nonalcoholic steatohepatitis (NASH) treatment. Although their glucose-lowering and insulin-sensitizing effects have been largely unraveled, the mechanisms by which they alleviate liver injury have only been scarcely addressed. Here, we aimed to unveil the mechanisms underlying the protective effects of FGF21 on NASH using APOE*3-Leiden.CETP mice, a well-established model for human-like metabolic diseases. Liver-specific FGF21 overexpression was achieved in mice, followed by administration of a high-fat high-cholesterol diet for 23 weeks. FGF21 prevented hepatic lipotoxicity, accompanied by activation of thermogenic tissues and attenuation of adipose tissue inflammation, improvement of hyperglycemia and hypertriglyceridemia, and upregulation of hepatic programs involved in fatty acid oxidation and cholesterol removal. Furthermore, FGF21 inhibited hepatic inflammation, as evidenced by reduced Kupffer cell (KC) activation, diminished monocyte infiltration and lowered accumulation of monocyte-derived macrophages. Moreover, FGF21 decreased lipid- and scar-associated macrophages, which correlated with less hepatic fibrosis as demonstrated by reduced collagen accumulation. Collectively, hepatic FGF21 overexpression limits hepatic lipotoxicity, inflammation and fibrogenesis. Mechanistically, FGF21 blocks hepatic lipid influx and accumulation through combined endocrine and autocrine signaling, respectively, which prevents KC activation and lowers the presence of lipid- and scar-associated macrophages to inhibit fibrogenesis.

## Introduction

The liver is the nexus of many metabolic pathways, including those of glucose, fatty acids (FAs) and cholesterol. In health, these metabolites are distributed to peripheral tissues while preventing long-lasting accumulation in the liver. In a pathological state, however, lipids may accrue in the liver, thereby impairing liver function and carving the path towards the development of nonalcoholic fatty liver disease (NAFLD) (1). NAFLD is considered a spectrum of liver diseases ranging from liver steatosis, characterized by lipid accumulation in hepatocytes, to nonalcoholic steatohepatitis (NASH) with hepatic steatosis, lobular inflammation, hepatocyte ballooning and varying degrees of fibrosis (2, 3). Patients diagnosed with NASH are predisposed to developing cirrhosis and hepatocellular carcinoma, among whom patients with severe liver fibrosis are at greatest risk of overall- and liver-related mortality (4). Despite this, there are currently no approved pharmaceutical therapeutics for NASH. Instead, lifestyle modifications remain the first-line treatment for NASH, although this is rarely attainable in the long term, and liver transplantation is still the sole intervention to treat the end-stage of NASH (2, 5). Thus, there is an unmet need for therapeutic strategies that control the progression of NASH, in particular of liver fibrosis, and reverse the underlying pathophysiology.

Current hypotheses suggest that adipose tissue dysfunction and lipid spillover leads to hepatic lipotoxicity, and thereby the initiation of NASH (6, 7), which further progresses through the inflammatory response triggered by hepatic lipotoxicity (7). This inflammatory response and subsequent fibrogenesis are primarily initiated by liver macrophages (8). Hepatic macrophages mainly consist of embryonically-derived macrophages, termed resident Kupffer cells (ResKCs), and monocyte-derived macrophages (MoDMacs) that are recruited from the circulation (9). In the steady state, ResKCs serve as sentinels for liver homeostasis. In NASH, liver injury caused by excess lipids and hepatocyte damage/death, triggers ResKC activation, leading to pro-inflammatory cytokine and chemokine release (10). This fosters the infiltration of newly-recruited monocytes into the liver, which gives rise to various pro-inflammatory and pro-fibrotic macrophage subsets (8, 10). Interestingly, recent preclinical and clinical studies have reported that modulation of ResKC activation, monocyte recruitment or macrophage differentiation, to some extent, can attenuate NASH (8, 11). In light of these findings, FGF21, a hepatokine with both lipid-lowering and anti-inflammatory properties (12, 13), has been brought to the foreground as a promising potential therapeutic to treat NASH.

The specificity of FGF21 action for various metabolic tissues is determined by the FGF receptor (FGFR) which forms a heterodimer with the transmembrane co-receptor β-Klotho (KLB) (14, 15). While the FGFR is ubiquitously expressed, KLB is primarily expressed in the liver and adipose tissue (14, 15), therefore possibly limiting FGF21 action to these tissues. Physiologically, FGF21 is considered a stress-induced hormone whose levels rise in metabolically compromised states, such as obesity (16) and NASH (17). The increased FGF21 in these pathologies is likely induced by an accumulation of lipids in the liver (18). As such, plasma FGF21 also positively correlates with the severity of steatohepatitis and fibrosis in patients with NASH (17). Induction of FGF21 is thought to mediate a compensatory response to limit metabolic dysregulation (19), although this level is not sufficient. Interestingly, two phase 2a clinical trials reported that pharmacological FGF21 treatment improves liver steatosis in NASH patients (20, 21). While an *in vivo* study testing the therapeutic potency of FGF21 in choline-deficient and high-fat diet-induced NASH has previously reported both anti-inflammatory and anti-fibrotic effects (22), detailed mechanistic understanding is still lacking.

In the present study, we aimed to elucidate the mechanisms underlying FGF21-mediated improvement of NASH, in particular of steatohepatitis and fibrogenesis. To this end, we used APOE*3-Leiden.CETP mice, a well-established model for human cardiometabolic diseases. These mice exhibit human-like lipoprotein metabolism, develop hyperlipidemia, obesity and inflammation when fed a high-fat high-cholesterol diet (HFCD), and develop fibrotic NASH closely resembling clinical features that accompany NASH in humans (23, 24). Moreover, these mice show human-like responses to both lipid-lowering and anti-inflammatory therapeutics during the development of metabolic syndrome (25–28). Here, we show that specific overexpression of FGF21 in the liver, resulting in increases circulating FGF21 levels, activates hepatic signaling associated with FA oxidation and cholesterol removal. In parallel, FGF21 activates thermogenic tissues and reduces adipose tissue inflammation, thereby protecting against adipose tissue dysfunction, hyperglycemia and hypertriglyceridemia. As a consequence, FGF21 largely limits lipid accumulation in the liver and potently blocks hepatic KC activation and monocyte recruitment, thereby preventing the accumulation of pro-inflammatory macrophages in the liver. In addition, FGF21 reduced the number of pro-fibrotic macrophages in the injured liver, potentially explaining why FGF21 counteracts all features of NASH, including hepatic steatosis, inflammation and fibrogenesis.

## Results

### Liver-specific FGF21 overexpression increases circulating FGF21 levels and protects against HFCD-induced body fat mass gain

We aimed to elucidate the underlying mechanisms of FGF21-mediated hepatoprotective effects on NASH, by using APOE*3-Leiden.CETP mice fed with a HFCD, a model that induces all stages of NASH in a human-like fashion and recapitulates the ultrastructural changes observed in NASH patients (23, 24). Since the liver is the main contributor to circulating FGF21 (14), we employed an adeno-associated virus vector 8 (AAV8) vector expressing codon-optimized FGF21 to induce liver-specific FGF21 overexpression in APOE*3-Leiden.CETP mice. Mice treated with either AAV8-FGF21 or AAV8-null as controls were fed with a HFCD for 23 weeks (**Figure 1A**). We confirmed liver-specific FGF21 overexpression by a large increase in *Fgf21* expression in the liver but not in adipose tissue, resulting in high circulating FGF21 levels that persisted throughout the study (**Figure 1B**). HFCD progressively and profoundly increased body weight over the experimental period, accompanied by increased white adipose tissue (WAT) and brown adipose tissue (BAT) weights relative to those of low fat low cholesterol (LFCD)-fed mice (**Figure 1C,D**). In favorable contrast, FGF21 reduced body weight in the first 3 weeks, after which body weight stabilized and remained lower than that of LFCD- and HFCD-fed mice by the end of the study (−18% and −35%, respectively; **Figure 1C**). Concomitantly, FGF21 decreased weights of gonadal WAT (gWAT; −67%), subcutaneous WAT (sWAT; −55%), interscapular BAT (iBAT; −41%) and subscapular BAT (−41%) to levels comparable to those observed in LFCD-fed mice (**Figure 1D**). These findings thus highlight the potent effects of FGF21 on preventing fat mass gain under NASH-inducing dietary conditions.

**Figure 1.**
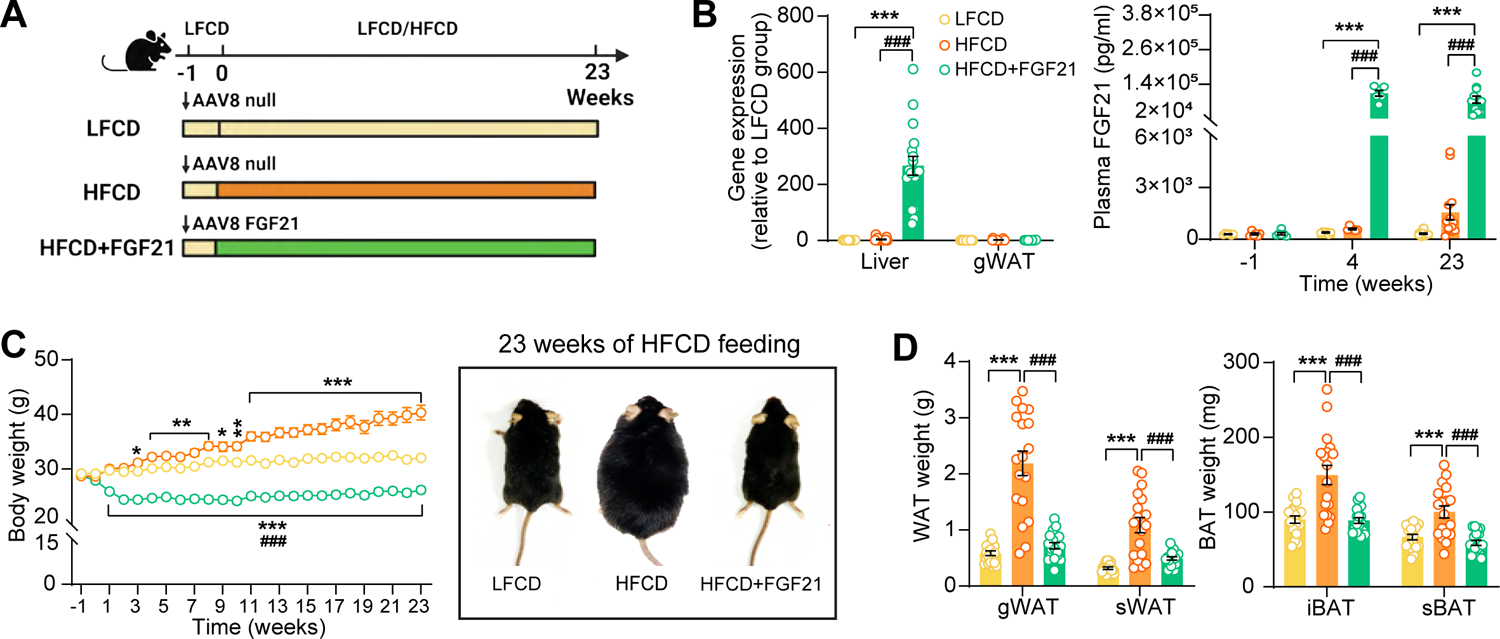
Liver-specific FGF21 overexpression increases circulating FGF21 levels and protects against HFCD-induced body fat mass gain. (A) Experimental set up. (B) At week 23, FGF21 mRNA expression in the liver and gWAT was quantified (n = 16-18). Plasma FGF21 levels were measured before (at week −1; pooled samples, n = 6 per group) and after (at week 4, pooled samples, n = 6 per group; week 23, n = 12-16 per group) AAV8-FGF21 administration. (C) Body weight was monitored throughout the experimental period (n = 17-18). (D) At week 23, brown adipose tissue (BAT) and white adipose tissue (WAT) depots were isolated and weighed (n = 18). Data are shown as mean ± SEM. Differences were assessed using one-way ANOVA followed by a Tukey post-test. \**P* < 0.05; \*\**P* < 0.01, \*\*\**P* < 0.001, compared with the LFCD group. ^###^*P* < 0.001, compared with the HFCD group. AAV8, adeno-associated virus 8; FGF21, fibroblast growth factor 21; gWAT, gonadal WAT; HFCD, high fat and high cholesterol diet; iBAT, interscapular BAT; LFCD, low fat and low cholesterol diet; sBAT, subscapular BAT; sWAT, subcutaneous white adipose tissue.

### FGF21 protects against HFCD-induced adipose tissue dysfunction

The profound fat mass-lowering effects of liver-derived FGF21 prompted us to examine its role in adipose tissue function. Since we and others have previously shown that FGF21 activates thermogenic adipose tissues (29, 30), we first performed histological analyses of BAT and sWAT, the adipose tissue that is most prone to browning (31). We observed that FGF21 prevented the HFCD-induced lipid overload in BAT (−66%) and increased uncoupling protein-1 (UCP-1) expression compared with both the LFCD- and HFCD-fed groups (+15% and +26%, respectively) (**Figure 2A**). In sWAT, FGF21 prevented HFCD-induced adipocyte hypertrophy (−41%), and increased the UCP-1 content (+94%) (**Figure 2B**). Among the adipose tissue depots, gWAT is most prone to diet-induced inflammation, and surgical removal of inflamed gWAT attenuates NASH in obese mice (32). Similar to sWAT, FGF21 protected against HFCD-induced adipocyte enlargement (−52%) in gWAT and in addition fully prevented the formation of crown-like structures (CLSs; −93%) (**Figure 2C**). In agreement with these findings, FGF21 suppressed the HFCD-induced expression of adhesion G protein-coupled receptor E1 (*Adgre1*; −56%), encoding the macrophage surface marker F4/80, in addition to decreased expression of the pro-inflammatory mediators tumor necrosis factor α (*Tnfa*; −60%), interleukin-1β (*Il1b*; −50%) and monocyte attractant chemokine C–C motif ligand 2 (*Ccl2*; −60%) (**Figure 2D**). Besides, FGF21 tended to upregulate *Klb* (+33%) and *Fgfr1* (+ 30%) expression compared to HFCD-fed mice (**Figure 2-figure supplement 1**). Moreover, consistent with the critical role of adiponectin in mediating the therapeutic benefits of FGF21 in adipose tissue(22, 33), FGF21 increased plasma adiponectin levels compared to both LFCD- and HFCD-fed mice (+93% and +133%, respectively; **Figure 2E**). These combined findings thus indicate that FGF21 prevents HFCD-induced adipose tissue dysfunction during NASH development.

**Figure 2.**
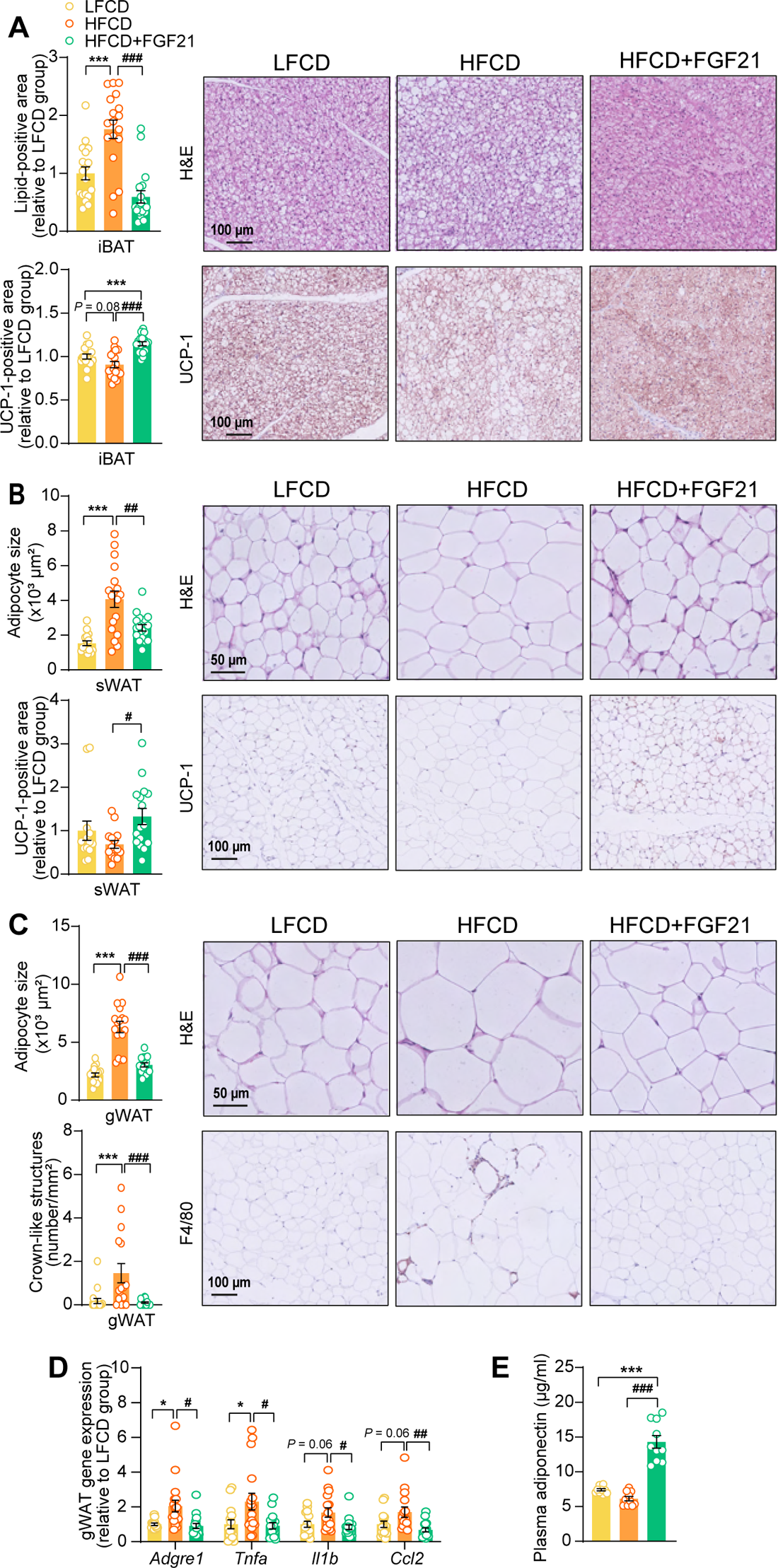
FGF21 protects against HFCD-induced adipose tissue dysfunction. (**A**) In iBAT, the lipid content and expression of uncoupling protein-1 (UCP-1) were quantified after H&E staining and UCP-1 immunostaining, respectively. (**B**) In sWAT, the adipocyte enlargement was assessed by H&E staining, and the tissue browning was evaluated by UCP-1 immunostaining. (**C**) In gWAT, the adipocyte hypertrophy was detected, and the number of CLSs was assessed, and (**D**) mRNA expression of pro-inflammatory markers was quantified. (**E**) Plasma adiponectin concentration in fasted blood plasma was measured at week 22. (**A**)-(**D**), n = 14-18 per group; (**E**), n = 10 per group. Differences were assessed using one-way ANOVA followed by a Tukey post-test. \**P* < 0.05, \*\*\**P* < 0.001, compared with the LFCD group. ^#^*P* < 0.05, ^##^*P* < 0.01, ^###^*P* < 0.001, compared with the HFCD group. *Adgre1*, adhesion G protein-coupled receptor E1; *Tnfa*, tumor necrosis factor α; *Il1b*, interleukin-1β; *Ccl2*, chemokine C–C motif ligand 2.

### FGF21 alleviates HFCD-induced hyperglycemia and hypertriglyceridemia

We next examined whether FGF21 confers its glucose and lipid lowering effects during NASH development. While HFCD induced hyperglycemia as compared to LFCD, FGF21 normalized fasting plasma glucose compared to LFCD, which was accompanied by lower glucose excursion after an intraperitoneal glucose tolerance test (**Figure 3A,B**). In addition, FGF21 normalized the plasma insulin and Homeostatic Model Assessment for Insulin Resistance index (**Figure 3C**), indicating that FGF21 restores insulin sensitivity to that observed in LFCD-fed mice. FGF21 did not prevent the HFCD-induced increase of plasma total cholesterol (TC) levels (**Figure 3-figure supplement 1A**), nor the distribution of cholesterol over the various lipoproteins (**Figure 3-figure supplement 1B**). Nonetheless, FGF21 strongly and consistently reduced fasting plasma triglyceride (TG) levels throughout the experimental period compared with LFCD- and HFCD-fed mice (−67% and −58%; at week 22), which was specific for very-low density lipoprotein (VLDL) and low density lipoprotein (LDL) (**Figure 3D**). In addition, an oral lipid tolerance test revealed that FGF21 prevented HFCD-induced lipid intolerance (**Figure 3E**). Taken together, FGF21 prevents the HFCD-induced increase in circulating glucose and reduces circulating TG levels beyond those observed in LFCD-fed mice.

**Figure 3.**
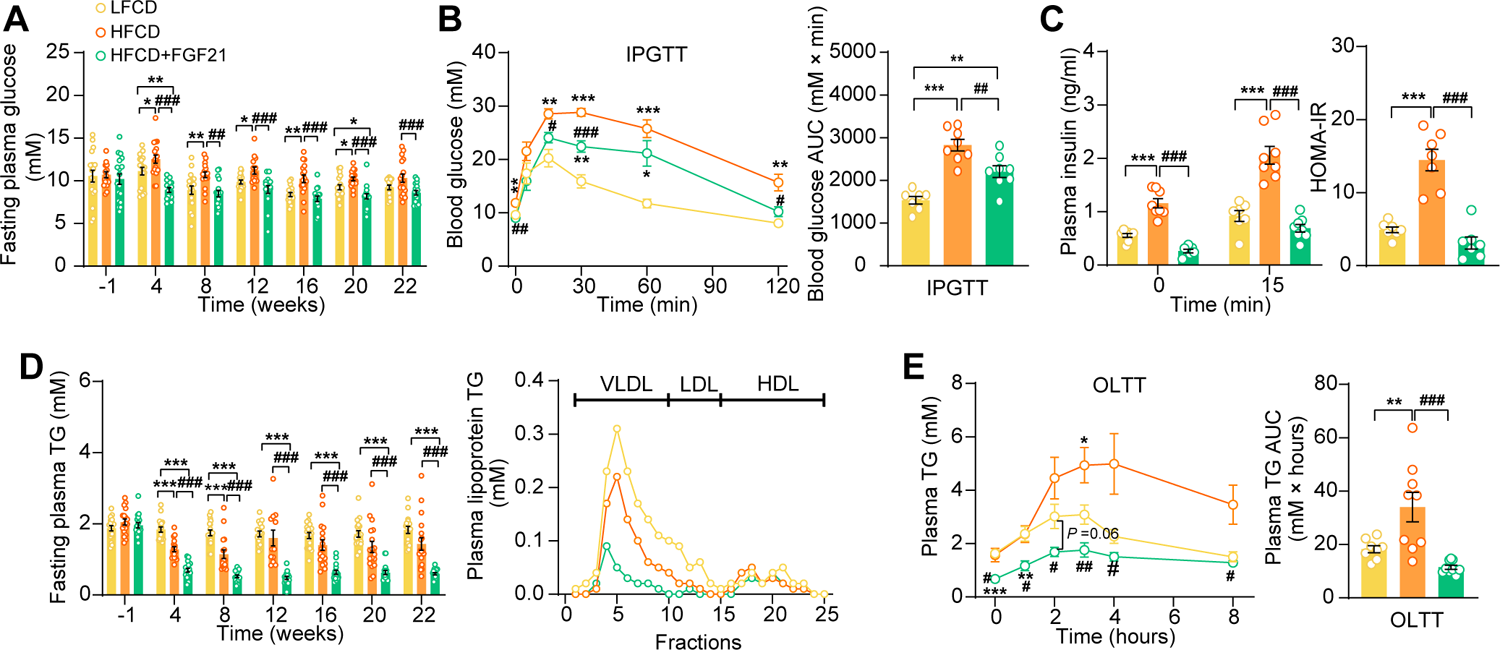
FGF21 alleviates HFCD-induced hyperglycemia and hypertriglyceridemia. (**A**) Fasting plasma glucose levels were measured during the experimental period. (**B**) At week 16, an intraperitoneal glucose tolerance test (IPGTT) was initiated. (**B**) The area under the curve (AUC) of plasma glucose during the IPGTT and (**C**) plasma insulin concentration in response to the IPGTT was determined at the indicated timepoints. (**C**) Homeostasis model assessment of insulin resistance (HOMA-IR) was determined from fasting glucose and insulin levels. (**D**) Fasting plasma TG levels were measured throughout the study. The distribution of triglyceride over lipoproteins was determined (pooled samples; n = 5 per group) from plasma of week 22. (**E**) At week 20, an oral lipid tolerance test (OLTT) was initiated, and AUC of plasma TG during the OLTT was calculated. (**A and D**), n = 14-18 per group; (**B-C**), n =7-8 per group; (**E**), n = 6-9 per group. Data are shown as mean ± SEM. Differences were assessed using one-way ANOVA followed by a Tukey post-test. \**P* < 0.05, \*\**P* < 0.01, \*\*\**P* < 0.001, compared with the LFCD group. ^#^*P* < 0.05, ^##^*P* < 0.01, ^###^*P* < 0.001, compared with the HFCD group.

### FGF21 protects against HFCD-induced hepatic steatosis, inflammation, and fibrogenesis

Then, we investigated the effects of FGF21 on liver steatosis, inflammation and fibrosis. FGF21 not only prevented HFCD-induced liver weight gain (−58%), but even reduced liver weight to a level lower than that of LFCD-fed mice (−40%; **Figure 4A,F**). Moreover, FGF21 abolished the HFCD-induced increase in steatosis, lobular inflammation and hepatocellular ballooning (**Figure 4B, Figure 4-figure supplement 1A,B**). Therefore, FGF21 completely prevented the HFCD-induced large increase in the NAFLD activity score (−74%; **Figure 4C,F**). Furthermore, FGF21 prevented collagen accumulation in the liver as assessed by Picrosirius Red staining (−58%; **Figure 4D,F**). We then measured hepatic concentration of hydroxyproline, a major constituent of collagen and thus a marker of extracellular matrix accumulation. In line with the hepatic collagen content, HFCD feeding increased the hepatic hydroxyproline content, which was prevented by FGF21 (−49%; **Figure 4E**). Taken together, our data demonstrate that FGF21 protects against HFCD-induced hepatosteatosis, steatohepatitis as well as fibrogenesis.

**Figure 4.**
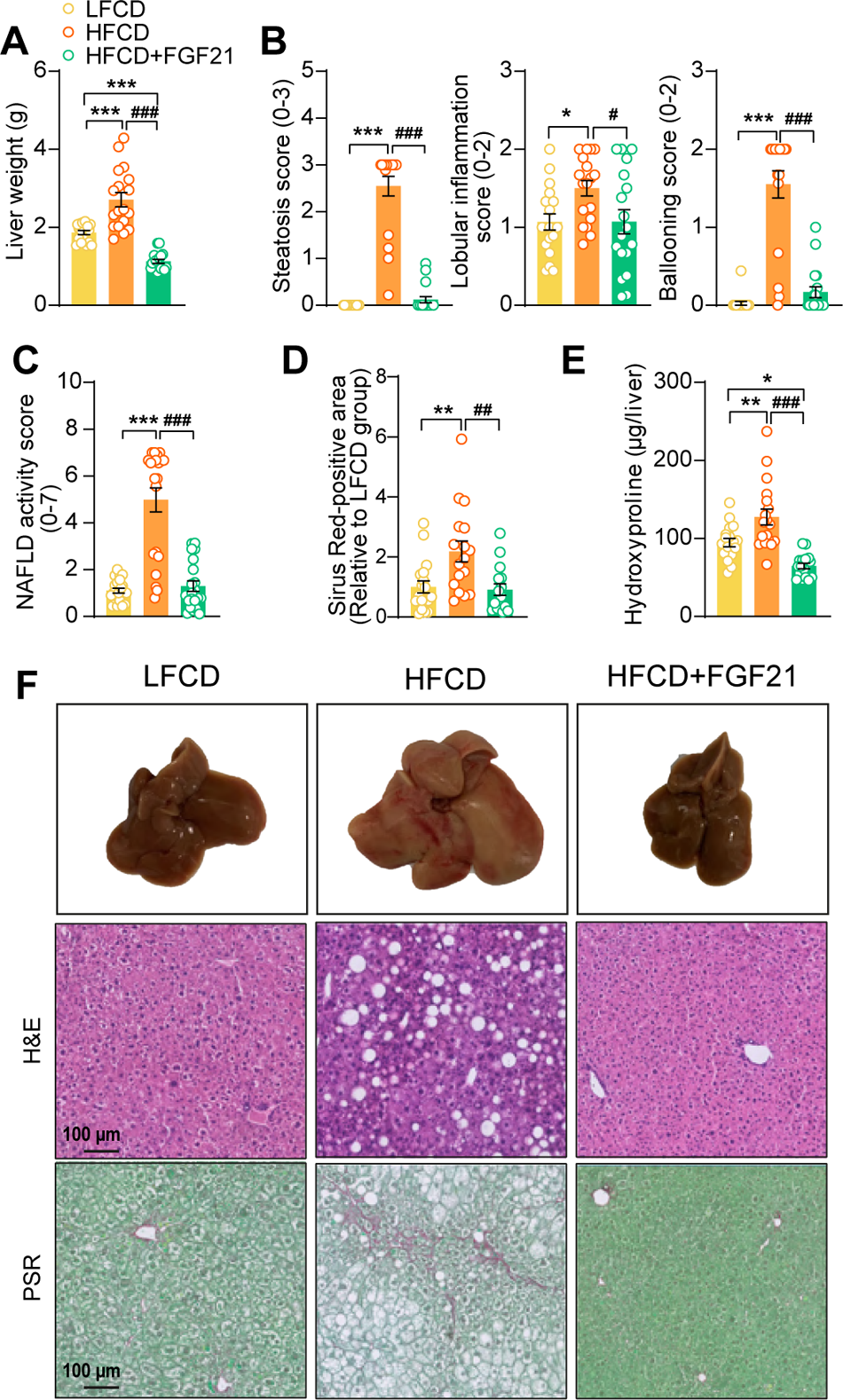
FGF21 protects against HFCD-induced hepatic steatosis, inflammation and fibrosis. (A) At week 23, liver weight was determined, and (**B**) scoring of histological features of steatosis, lobular inflammation and ballooning as well as (**C**) NAFLD activity was evaluated by H&E staining. (**D**) Liver fibrosis was assessed by Picrosirius Red (PSR) staining, and (**E**) hepatic hydroxyproline levels were determined. (**F**) Representative macroscopic, H&E and PSR pictures are shown. Data are shown as mean ± SEM (n = 16-18 per group). Differences were assessed using one-way ANOVA followed by a Tukey post-test. \**P* < 0.05; \*\**P* < 0.01, \*\*\**P* < 0.001, compared with the LFCD group. ^##^*P* < 0.01; ^###^*P* < 0.001, compared with the HFCD group.

### FGF21 abolishes liver lipotoxicity, accompanied by activation of hepatic signaling involved in FA oxidation and cholesterol removal

In the context of NASH, pro-inflammatory responses and fibrogenesis occur when hepatocytes are injured by lipotoxicity (7, 34). Indeed, 23 weeks of HFCD feeding promoted aberrant accumulation of TG as well as TC in the liver (**Figure 5A**). In agreement with the data presented in **Figure 4**, FGF21 abrogated the HFCD-induced increase in hepatic TG levels (−62%) and tended to decrease hepatic TC levels (−22%), resulting in smaller lipid droplets (**Figure 5A**). In addition to reduced lipid overflow from WAT, we reasoned that FGF21 may also directly act on the liver to prevent HFCD-induced liver lipotoxicity. In agreement, compared to both LFCD- and HFCD-fed mice, FGF21 profoundly upregulated the expression of *Klb* (+150% and +223%), *Fgfr1* (+57% and +79%), *Fgfr2* (+97% and +77%), and *Fgfr4* (+53% and +67%) (**Figure 5-figure supplement 1**). We next quantified the hepatic expression of key genes involved in FA and cholesterol handling. FGF21 did not attenuate the HFCD-induced increased expression of FA translocase cluster of differentiation 36 (C*d36*) (**Figure 5-supplement 2A**). In favorable contrast, compared to both LFCD- and HFCD-fed mice, FGF21 did increase the expression of carnitine palmitoyl transferase 1α (*Cpt1a*, +66% and +53%), peroxisome proliferator-activated receptor α (*Ppara*, +67% and +53%) and peroxisome proliferator-activated receptor γ coactivator 1α (*Pgc1a*; +188% and +225%), all of those genes being key players involved in FA oxidation (**Figure 5B**). Moreover, compared to LFCD- and HFCD-fed mice, FGF21 increased the expression of apolipoprotein B (*Apob*, +26% and +38%), which is involved in VLDL secretion (**Figure 5-figure supplement 2B**). Furthermore, FGF21 upregulated the expression of ATP-binding cassette transporter G member 5 (*Abcg5*; 7-fold and 2-fold), crucial for biliary secretion of neutral sterols (**Figure 5C**), increased the expression of cholesterol 7α-hydroxylase (*Cyp7a1*; +94% and +109%), a key gene involved in the classic bile acid synthesis pathway (**Figure 5D**), and restored the expression of sterol 27-hydroxylase (+38%), involved in the alternative bile acid pathway (**Figure 5D**). Considering that bile acid synthesis is a major pathway for hepatic cholesterol disposal (35), FGF21 likely regulates bile acid metabolism to prevent HFCD-induced cholesterol accumulation in the liver. Collectively, our data indicate that FGF21 increases the hepatic expression of key genes involved in β-oxidation and cholesterol removal, which together with reduced lipid overload from WAT may explain FGF21-induced alleviation of liver lipotoxicity under NASH-inducing dietary conditions.

**Figure 5.**
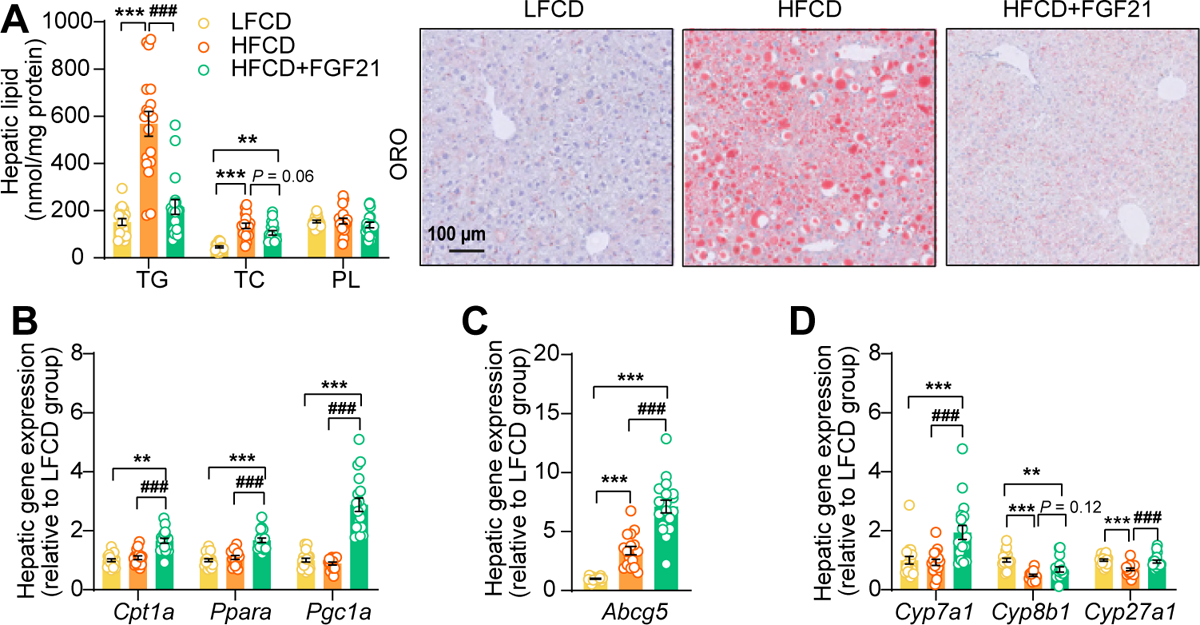
FGF21 abolishes liver lipotoxicity, accompanied by activation of hepatic signaling involved in FA oxidation and cholesterol removal. (**A**) Triglyceride (TG), total cholesterol (TC) and phospholipid (PL) levels were determined in the liver (n = 18 per group), and representative Oil Red O (ORO) pictures are shown. (**B**) The relative mRNA expression of genes involved in fatty acid oxidation and (**C and D**) cholesterol removal (n = 15-18 per group) were determined in the liver. Data are shown as mean ± SEM. Differences were assessed using one-way ANOVA followed by a Tukey post-test. \*\**P* < 0.01, \*\*\**P* < 0.001, compared with the LFCD group. ^###^*P* < 0.001, compared with the HFCD group. *Abcg5*, ATP-binding cassette transporter G member 5; *Cpt1a*, carnitine palmitoyl transferase 1α; *Cyp7a1*, cholesterol 7α-hydroxylase; *Cyp8b1*, sterol 12α-hydroxylase; *Cyp27a1*, sterol 27-hydroxylase; *Pgc1a*, peroxisome proliferator-activated receptor gamma coactivator 1α; *Ppara*, peroxisome proliferator-activated receptor α.

### FGF21 prevents activation of various KC subsets

Then, we performed an in-depth phenotyping of hepatic immune cells using spectral flow cytometry. For this, we developed a panel that identifies most major immune cell subsets (for gating strategy see **Figure 6-figure supplement 1A**). As compared to LFCD, HFCD tended to reduce total CD45^+^ leukocytes, which were increased by FGF21 (**Figure 6-figure supplement 1B**). Combining conventional gating and dimension-reduction analysis through uniform manifold approximation and projection allowed to identify FGF21-induced changes in cell subset abundance (**Figure 6A**). FGF21 prevented HFCD-induced loss of eosinophils, neutrophils and B cells, and increased numbers of dendritic cells and T cells compared with those observed in both LFCD- and HFCD-fed mice (**Figure 6-figure supplement 1B**). More importantly, FGF21 increased the number of total KCs compared with that of both LFCD- and HFCD-fed mice (+63% and +156; **Figure 6-figure supplement 1B**), attenuated HFCD-induced monocyte recruitment (−18%), and tended to repress the HFCD-induced increase in hepatic MoDMacs (−42%; **Figure 6-figure supplement 1B**).

**Figure 6.**
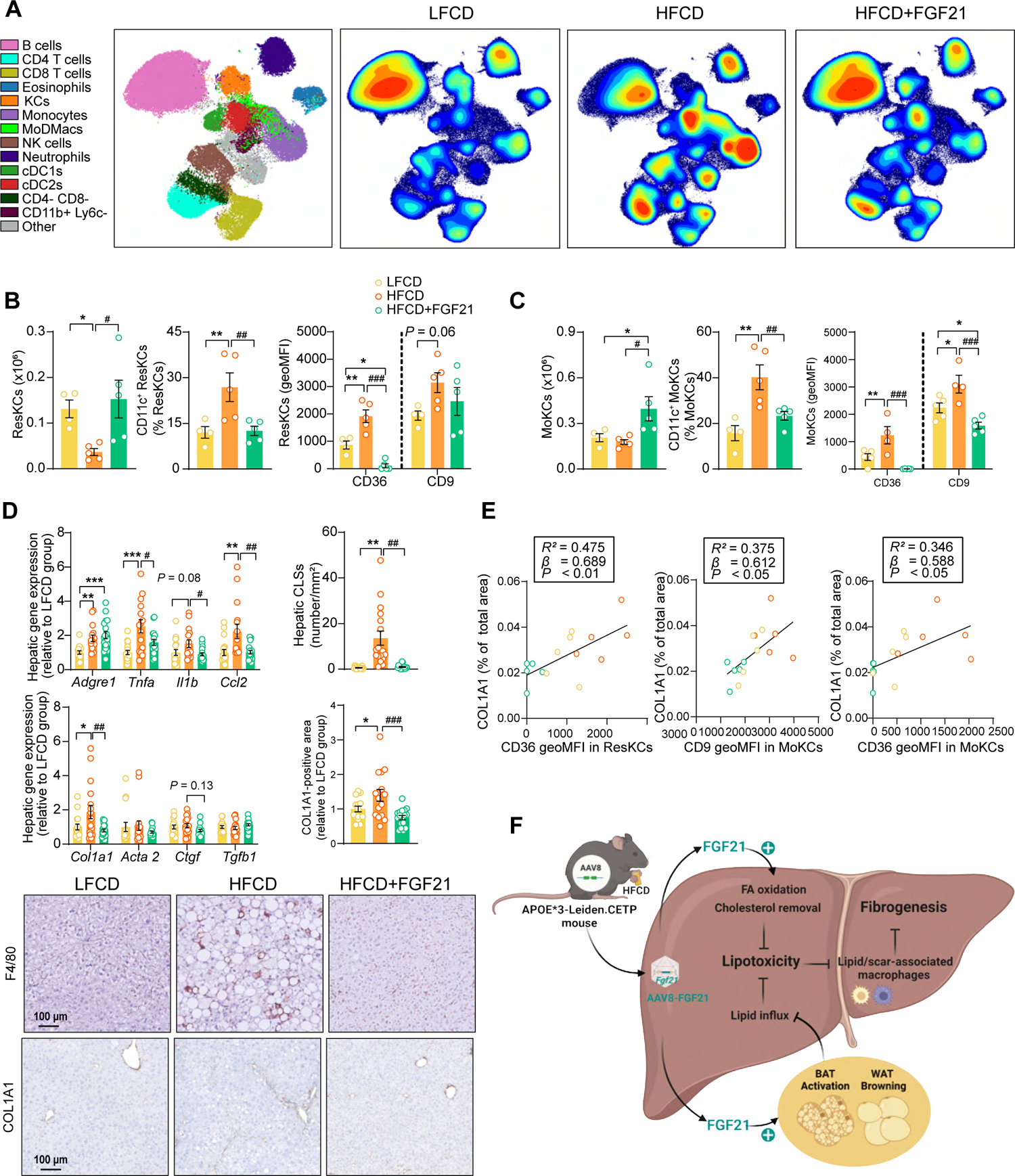
FGF21 modulates hepatic macrophage pool and protects against COL1A1 accumulation, as predicted by the reduction of CD36^hi^ KCs and CD9^hi^ KCs. (**A**) Uniform manifold approximation and projection for dimension reduction (UMAP) of immune cell subsets from livers after 23-week of intervention. (**B**) The number of resident KCs (ResKCs), the proportion of CD11c^+^ ResKCs, and the expression of CD36 and CD9 in ResKCs were quantified. (**C**) The amount of monocyte-derived KCs (MoKCs) was assessed, the percentage of CD11c^+^ MoKCs was determined, the CD36 and CD9 expression levels in MoKCs were quantified. (**D**) Hepatic inflammation was evaluated by pro-inflammatory gene expression and the formation of CLSs within the liver. The mRNA expression of liver fibrogenesis markers was quantified, and the protein expression of collagen type 1α 1 (COL1A1) was determined. (**E**) The expression of CD36 in ResKCs, and the expression of CD9 and CD36 in MoKCs were plotted against COL1A1-positive area in the liver. (**F**) Mechanistic model. Data are shown as mean ± SEM (**A-B and E**, n = 4-5 per group; **D**, n = 16-18 per group). Linear regression analyses were performed. Differences were assessed using one-way ANOVA followed by a Fisher’s LSD test. \**P* < 0.05, \*\**P* < 0.01, \*\*\**P* < 0.001, compared with the LFCD group. ^#^*P* < 0.05, ^##^*P* < 0.01, ^###^*P* < 0.001, compared with the HFCD group*. Acta2*, actin α 2; *Ctgf*, connective tissue growth factor; FA, fatty acid; *Tgfb1*, transforming growth factor-β.

During the development of NASH, MoDMacs can gradually seed in KC pool by acquiring ResKCs identity and replacing the dying ResKCs (36). These recruited MoKCs can have both detrimental and supportive roles, contributing to increase in pathology during fibrosis onset, but hastening recovery when the damage-evoking agent is attenuated/removed (37). In light of this, we assessed the abundance and phenotype of ResKCs and monocyte-derived KCs (MoKCs). We observed that FGF21 completely abolished the HFCD-induced reduction of the number of ResKCs (+319%) and potently protected against HFCD-induced ResKC activation as shown by decreased proportion of CD11c^+^ ResKCs (−53%; **Figure 6B**). FGF21 also completely abolished the HFCD-induced upregulation of CD36 in ResKCs, to levels that are even lower than those in LFCD-fed mice (−88% *vs.* LFCD; −94% *vs.* HFCD; **Figure 6B**). In addition, FGF21 increased the number of MoKCs compared with that of both LFCD- and HFCD-fed mice (+92% and +123%), and prevented the HFCD-induced increase in the abundance of CD11c^+^ MoKCs (−42%) (**Figure 6C**). Strikingly, compared to both LFCD- and HFCD-fed mice, FGF21 downregulated CD9 (−32% and −49%) and CD36 (−98% and −100%) in MoKCs (**Figure 6C**). Furthermore, FGF21 profoundly repressed HFCD-induced upregulation of hepatic *Tnfa* (−37%), *Il1b* (−41%) and *Ccl2* (−54%) expression to levels comparable to those in LFCD-fed mice (**Figure 6D)**, which is in line with the observation that FGF21 prevents KC activation. Given that CD36^hi^ ResKCs and CD36^hi^/ CD9^hi^ MoKCs are involved in the formation of hepatic CLSs(10, 37–39), we next assessed CLSs and observed that FGF21 completely prevented the HFCD-induced formation of CLSs in the liver (−93%; **Figure 6D**). These data demonstrate that FGF21 inhibits the activation of ResKCs and MoKCs and prevents the accumulation of CD36^hi^ ResKCs and CD36^hi^/ CD9^hi^ MoKCs under dietary conditions that result in NASH, which likely contribute to the beneficial effects of FGF21 on hepatic inflammation and fibrosis.

### FGF21 protects against COL1A1 accumulation, as predicted by the reduction of CD36^hi^ KCs and CD9^hi^ KCs

To further evaluate whether FGF21-induced reductions of lipid-associated macrophages (i.e., CD36^hi^ ResKCs and CD36^hi^ MoKCs) (38) and scar-associated macrophages (i.e., CD9^hi^ MoKCs) (40), are implicated in fibrogenesis, we performed multiple univariate regression analyses. These revealed that both NAFLD activity and liver fibrosis were associated with both CD36^hi^ ResKCs, CD36^hi^ MoKCs and CD9^hi^ MoKCs (**Figure 6-figure supplement 2A-D**), indicating that FGF21 likely improves liver fibrosis by reducing these lipid- and scar-associated macrophages. To further understand the underlying mechanisms by which FGF21 prevents liver fibrosis, we measured hepatic expression of key genes involved in fibrogenesis (**Figure 6D**). FGF21 tended to decrease the expression of connective tissue growth factor (*Ctgf*; −27%), a major fibrogenic factor, and normalized the HFCD-induced increased expression of its downstream target collagen type Iα 1 (*Col1a1*; −61%; **Figure 6D**). This finding was confirmed by immunohistochemistry, revealing that FGF21 reduced hepatic COL1A1 accumulation (−46%; **Figure 6D**). Furthermore, univariate regression analysis revealed that COL1A1 expression is predicted by CD36^hi^ ResKCs, CD36^hi^ MoKCs and CD9^hi^ MoKCs (**Figure 6E, Figure 6-figure supplement 2E**). Taken together, these data indicate that FGF21 reduces lipid- and scar-associated macrophages to inhibit COL1A1 synthesis and prevent fibrogenesis.

## Discussion

Several FGF21 analogues are currently being evaluated in clinical trials for the treatment of NASH (20, 21). While the protective effect of pharmacological intervention with long-acting FGF21 on human liver steatosis has been uncovered (20, 21, 41), mechanisms underlying attenuated steatosis as well all the anti-inflammatory and anti-fibrotic effects of FGF21 on NASH are still largely unexplored. Therefore, we set out to elucidate mechanisms by which FGF21 beneficially modulates these various aspects of NASH in HFCD-fed APOE*3-Leiden.CETP mice, a well-established model for diet-induced NASH (23, 24). Based on our findings, we propose that FGF21 attenuates liver lipotoxicity via endocrine signaling to adipose tissue to induce thermogenesis, thereby preventing adipose tissue dysfunction to reduce lipid overflow to the liver, as well as autocrine signaling to the liver to increase FA oxidation and cholesterol removal. In addition, FGF21 prevents KC activation, monocyte recruitment and the formation of lipid- and scar-associated macrophages, thereby likely inhibiting collagen accumulation and alleviating liver fibrogenesis.

Hepatic lipotoxicity is one of the major risk factors determining the progression of liver steatosis into NASH, as shown in multiple clinical studies with obese patients (42–44). By feeding APOE*3-Leiden.CETP mice a diet rich in fat and cholesterol, we mimicked a situation in which a positive energy balance induces many aspects of the metabolic syndrome, including insulin resistance, obesity with increased fat accumulation, and hepatic lipotoxicity indicated by hepatomegaly with aberrant accumulation of TG as well as TC. Hepatic lipotoxicity likely results from lipid overflow from insulin-resistant adipose tissue towards the liver in combination with hepatic insulin resistance that prevents insulin-stimulated outflow of lipids (45). Within this dietary context, we applied a single administration of an AAV8 vector encoding codon-optimized FGF21, which resulted in liver-specific FGF21 overexpression. Since the codon-optimized FGF21 mitigates the poor pharmacokinetic properties of native FGF21, including its short plasma half-life (0.5-2 hours) by reducing proteolytic degradation(45), an elevated level of circulating FGF21 was reached throughout the dietary intervention period. By this strategy, we mimicked the situation in which circulating FGF21 predominantly derives from the liver (46). Indeed, circulating FGF21 correlates well with the hepatic expression of FGF21 (47). Interestingly, hepatic expression of FGF21 fully prevented the diet-induced increase in liver weight, liver lipids (i.e., TG and TC) and steatosis score.

These lipotoxicity-protective effects of FGF21 can partially be explained by endocrine effects of liver-derived FGF21 on adipose tissue, which besides the liver has high expression of β-Klotho, the co-receptor of the FGFR (14, 15). Indeed, FGF21 fully prevented the HFCD-induced increase in weights of WAT and BAT, with decreased lipid accumulation in these adipose tissue depots as well as induction of BAT activation and WAT browning. These data imply that FGF21 greatly induces thermogenesis which highly increases energy expenditure, consistent with the thermogenic responses observed for recombinant FGF21 in mice fed with an obesogenic diet (29) or atherogenic diet (30). Activation of thermogenic tissues by classical β-adrenergic receptor largely increases the uptake of circulating lipoprotein-derived FAs by BAT and beige WAT (48), which we recently also demonstrated for recombinant FGF21 (30). This can thus at least partly explain the marked TG-lowering effect of FGF21 observed in the current study. Thermogenic activation also increases the uptake and combustion of glucose, although the glucose-lowering and insulin-sensitizing effects of FGF21 can also be explained by attenuated WAT inflammation in combination with increased adiponectin expression as well as improved liver insulin sensitivity (30, 33, 49).

Besides endocrine FGF21 signaling in adipose tissue, liver lipotoxicity is likely further prevented by autocrine FGF21 signaling. Indeed, we showed that liver-specific FGF21 overexpression increased hepatic expression of genes involved in FA oxidation (*Cpt1a*, *Ppara*, *Pgc1a*), biliary cholesterol secretion (*Abcg5*), bile acids synthesis (*Cyp7a1*) and VLDL production (*Apob*). Of note, these observations are in line with previous reports showing increased FA oxidation (50) and upregulated *Abcg5* (51), *Cyp7a1* (51, 52) and *Apob* (30) in the liver upon FGF21 treatment. Altogether, the marked protective effects of FGF21 on HFCD-induced hepatic lipotoxicity likely results from combined endocrine and autocrine signaling, leading to reduced lipid influx from adipose tissue to the liver coupled to the activation of hepatic FA oxidation and cholesterol elimination pathways. Our observations may likely explain the recent clinical findings that treatment with FGF21 analogues in patients with NASH not only reduced hepatic steatosis (20, 21) but also increased hepatic bile acid synthesis and further promoted cholesterol removal, lowering the risk for further hepatic lipotoxicity (53).

While NASH is initiated by hepatic lipotoxicity, NASH progression is mainly driven by impaired KC homeostasis and subsequent liver inflammation (54). Therefore, we investigated in depth the inflammatory response in the liver through a combination of immunohistochemistry, flow cytometry and gene expression analyses. HFCD feeding induced an array of inflammatory effects, including increased lobular inflammation, hepatocyte ballooning and NAFLD activity scores as well as increased inflammatory foci and CLSs, accompanied by a reduction in ResKCs with a relative increase in CD11c^+^ ResKCs, and an increase in MoDMacs and CD11c^+^ MoKCs. These observations are likely explained by lipotoxicity-related damage to ResKCs, and release of TNFα, IL-1β and MCP-1 (*Ccl2*), both activating various downstream pro-inflammatory mediators as well as promoting monocyte recruitment to remodel the KC pool(36, 55) and further exacerbating hepatic inflammation (10, 38, 54, 56, 57). Importantly, FGF21 prevented most of these HFCD-induced inflammatory responses, as it normalized lobular inflammation, hepatocyte ballooning and NAFLD activity scores and CLSs, and reduced pro-inflammatory activation of various KC subsets.

Fibrosis has been identified as the most important predictor of prognosis in NAFLD patients, and therefore a main target in experimental pharmacological approaches (58). HFCD feeding during 23 weeks induced early signs of fibrosis, as evident from an increased *Col1a1* expression and COL1A1 content, accompanied by an increased content of the hydroxyproline. Importantly, FGF21 blocked liver fibrogenesis, and decreased the hydroxyproline content. These alterations were accompanied with reductions in lipid-associated macrophages (i.e., CD36^hi^ ResKCs/MoKCs) (38) and scar-associated macrophages (i.e., CD9^hi^ MoKCs) (40). In fact, when analysing the mouse groups together, CD36^hi^ ResKCs/MoKCs and CD9^hi^ MoKCs positively correlated with liver fibrosis as reflected by hydroxyproline content and COL1A-positive area, suggesting that these lipid- and scar-associated macrophages are involved in fibrogenesis in our model. Indeed, high numbers of CD9^hi^ macrophages have been found in fibrotic regions of the liver (37, 39, 40, 55), and these cells are able to prime quiescent primary murine hepatic stellate cells to upregulate the expression of fibrillar collagen through CTGF (40), thereby promoting and exacerbating liver fibrosis. Therefore, we speculate that FGF21 protects against early liver fibrosis likely through preventing the accumulation of CD36^hi^/CD9^hi^ KCs, thereby inhibiting activation of hepatic stellate cells to produce collagen.

In conclusion, hepatic overexpression of FGF21 in APOE*3-Leiden.CETP mice limits diet-induced hepatic lipotoxicity, inflammation and fibrogenesis. Through a combination of endocrine and autocrine signaling, FGF21 reduces hepatic lipid influx and accumulation, respectively. This results in reduced macrophage activation and monocyte recruitment with less presence of lipid- and scar-associated macrophages, limiting activation of hepatic stellate cells to produce collagen (for graphic summary see **Figure 6F**). As such, our studies provide a mechanistic explanation for the hepatoprotective effects of FGF21 analogues in recent clinical trials including reduction in steatosis (20, 21, 53) as well as the fibrotic marker N-terminal type III collagen pro-peptide (20, 21), and further highlight the potential of FGF21 for clinical implementation as a therapeutic in the treatment of advanced NASH.

## Materials and Methods

Please see the **Supporting Information** for a detailed description of all experimental procedures.

### Animals and treatments

Male APOE*3-Leiden.CETP mice (on a C57BL/6J background) were generated as previously described (59). Mice at the age of 10-12 weeks were group-housed (2-4 mice per cage) under standard conditions (22°C, 12/12-hour light/dark cycle) with *ad libitum* access to water and a LFCD (Standard Rodent Diet 801203, Special Diets Services, United Kingdom), unless indicated otherwise. Then, based on body weight and 4-hour (9.00-13.00) fasted plasma glucose, TG and TC levels, these mice were randomized into three treatment groups (n = 18 per group), after which they received either AAV8-FGF21, a liver-tropic AAV8 capsid vector expressing FGF21 under the control of a liver specific apolipoprotein E/antitrypsin promoter (HFCD+FGF21 group; 2×10^10^ genome copies per mouse), or with the same genome copy number of AAV8-null (HFCD and LFCD groups) via a single intravenous injection. After one week of recovery, mice in the HFCD+FGF21 and HFCD groups were switched to a HFCD (60% fat and 1% cholesterol; C1090-60, Altromin, Germany) and maintained on the diet for 23 weeks. An intraperitoneal glucose tolerance test (n = 8 per group) and an oral lipid tolerance test (n = 10 per group) were performed at week 16 and week 20, respectively. Flow cytometry (n = 5 per group) was conducted at week 23.

### Statistics

Comparisons among three groups were analyzed using one-way ANOVA followed by a Tukey post-test, unless indicated otherwise. Data are presented as mean ± SEM, and a *P* value of less than 0.05 was considered statistically significant. All statistical analyses were performed with GraphPad Prism 9.01 for Windows (GraphPad Software Inc., California, CA, USA).

### Study approval

All animal experiments were carried out according to the Institute for Laboratory Animal Research Guide for the Care and Use of Laboratory Animals, and were approved by the National Committee for Animal Experiments (Protocol No. AVD1160020173305) and by the Ethics Committee on Animal Care and Experimentation of the Leiden University Medical Center (Protocol No. PE.18.034.041).

## Acknowledgments

This work was supported by the Dutch Diabetes Research Foundation (2015.81.1808 to M.R.B.); the Netherlands Organisation for Scientific Research-NWO (VENI grant 91617027 to Y.W.); Chinese Scholarship Council grants (CSC 201606010321 to E.Z.); the Novo Nordisk Foundation (NNF18OC0032394 to M.S.); and the Netherlands Cardiovascular Research Initiative: an initiative with support of the Dutch Heart Foundation (CVON-GENIUS-2 to P.C.N.R.). The authors also thank T.C.M. Streefland, A.C.M. Pronk, R.A. Lalai and H.C.M. Sips from Department of Medicine, the Division of Endocrinology, Leiden University Medical Center for technical assistance.

## Conflict of interest

ACA, AP, SO, MU, IA, YI, KW and XRP are employees of AstraZeneca.

## Data availability

All data generated or analyzed during this study are included in the manuscript and supporting file.

## Author contributions

CL designed the study, carried out the research, analyzed and interpreted the results, and wrote and revised the manuscript. MS interpreted the results, reviewed and revised the manuscript and obtained the funding. BS and EZ carried out the research and reviewed the manuscript. JML, HJPZ and BG designed and advised the study, interpreted the results and reviewed the manuscript. MET advised the study and reviewed the manuscript. ACA, SO and KW advised the study, interpreted the results and reviewed the manuscript. AP designed AAV8-FGF21 vectors and edited the manuscript. MU and IA analyzed and interpreted the results and reviewed the manuscript. YI and X-RP provided AAV8-FGF21 vectors, advised the study, interpreted the results and reviewed the manuscript. MRB advised the study and reviewed the manuscript. YW designed and advised the study, interpreted the results, reviewed and revised the manuscript. PCNR designed and advised the study, interpreted the results, edited, reviewed and revised the manuscript and obtained the funding.

## Figure supplements

**Figure 2-figure supplement 1.**
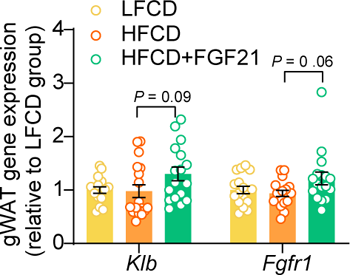
Liver-specific FGF21 overexpression tends to upregulate mRNA expression of FGF21 receptor 1 (FGFR1) and co-receptor β-Klotho (KLB) in white adipose tissue (WAT). The mRNA expression of KLB and FGFR1 in gonadal WAT (gWAT). Data are shown as mean ± SEM (n = 16-18 per group). Differences were assessed using one-way ANOVA followed by a Tukey post-test.

**Figure 3-figure supplement 1.**
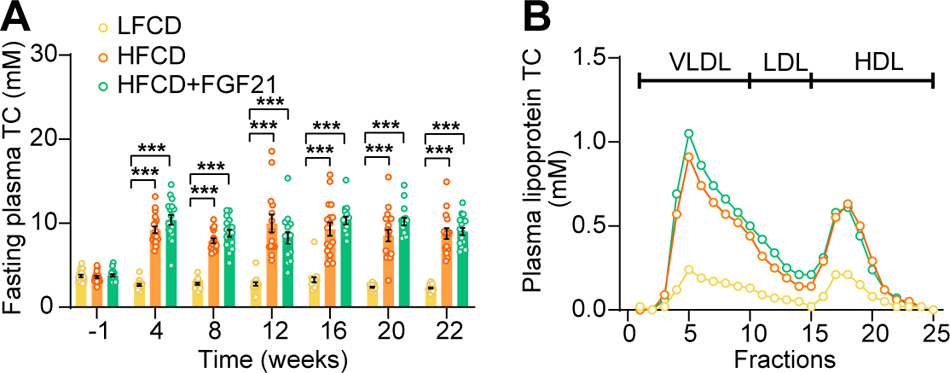
HFCD increases fasting cholesterol levels. (**A**) Fasting plasma total cholesterol (TC) levels were measured over a 23-week intervention period (n = 14-18 per group), and (B) the distribution of the cholesterol over circulating lipoproteins was assessed at week 22 (pooled samples; n = 18 per group). Data are shown as mean ± SEM. Differences were assessed using one-way ANOVA followed by a Tukey post-test. \*\*\**P* < 0.001, compared with the LFCD group. VLDL, very low-density lipoprotein; LDL, low-density lipoprotein; HDL, high-density lipoprotein.

**Figure 4-figure supplement 1.**
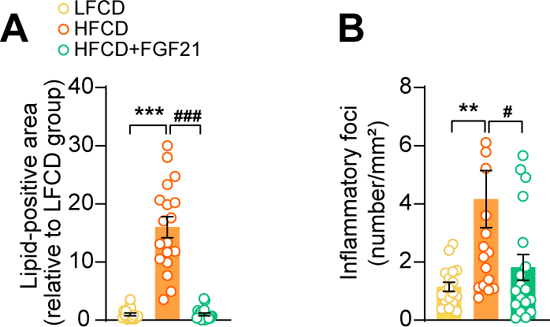
FGF21 abolishes HFCD-induced increase of hepatic lipid-positive area and the number of inflammatory foci. At week 23, (**A**) hepatic lipid droplet content and (**B**) inflammatory foci numbers were assessed by H&E staining. Data are shown as mean ± SEM (n = 18 per group). Differences were assessed using one-way ANOVA followed by a Tukey post-test. \*\**P* < 0.01, \*\*\**P* < 0.001, compared with the LFCD group. ^#^*P* < 0.01 ^###^*P* < 0.001, compared with the HFCD group.

**Figure 5-figure supplement 1.**
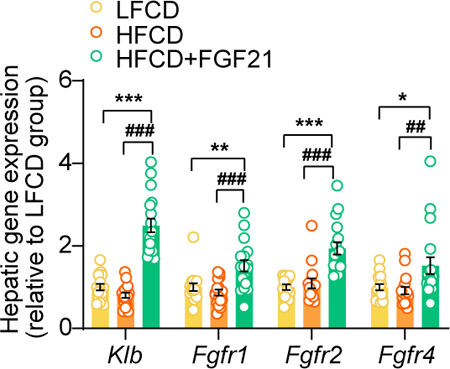
Liver-specific FGF21 overexpression upregulates hepatic mRNA expression of FGF21 receptors (FGFRs) and co-receptor β-Klotho (KLB). The mRNA levels of KLB and FGFRs in the liver. Data are shown as mean ± SEM (n = 14-18 per group). Differences were assessed using one-way ANOVA followed by a Tukey post-test. \**P* < 0.05, \*\**P* < 0.01, \*\*\**P* < 0.001, compared with the LFCD group. ^##^*P* < 0.01, ^###^*P* < 0.001, compared with the HFCD group.

**Figure 5-figure supplement 2.**
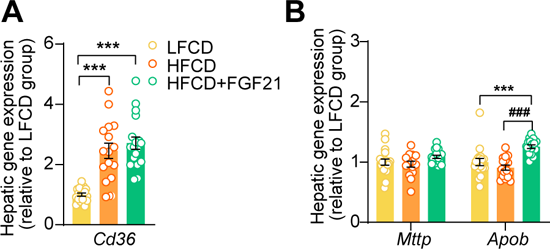
FGF21 increases apolipoprotein B mRNA (*Apob)* expression in the liver. At end of the study, hepatic expression of genes involved in (**A**) fatty acid uptake and (**B**) VLDL production was quantified (n = 15-18 per group). Data are shown as mean ± SEM. Differences were assessed using one-way ANOVA followed by a Tukey post-test. \*\*\**P* < 0.001, compared with the LFCD group. ^###^*P* < 0.001, compared with the HFCD group. *Apob*, apolipoprotein B*; Cd36*, cluster of differentiation 36; *Mttp*, microsomal triglyceride transfer protein.

**Figure 6-figure supplement 1.**
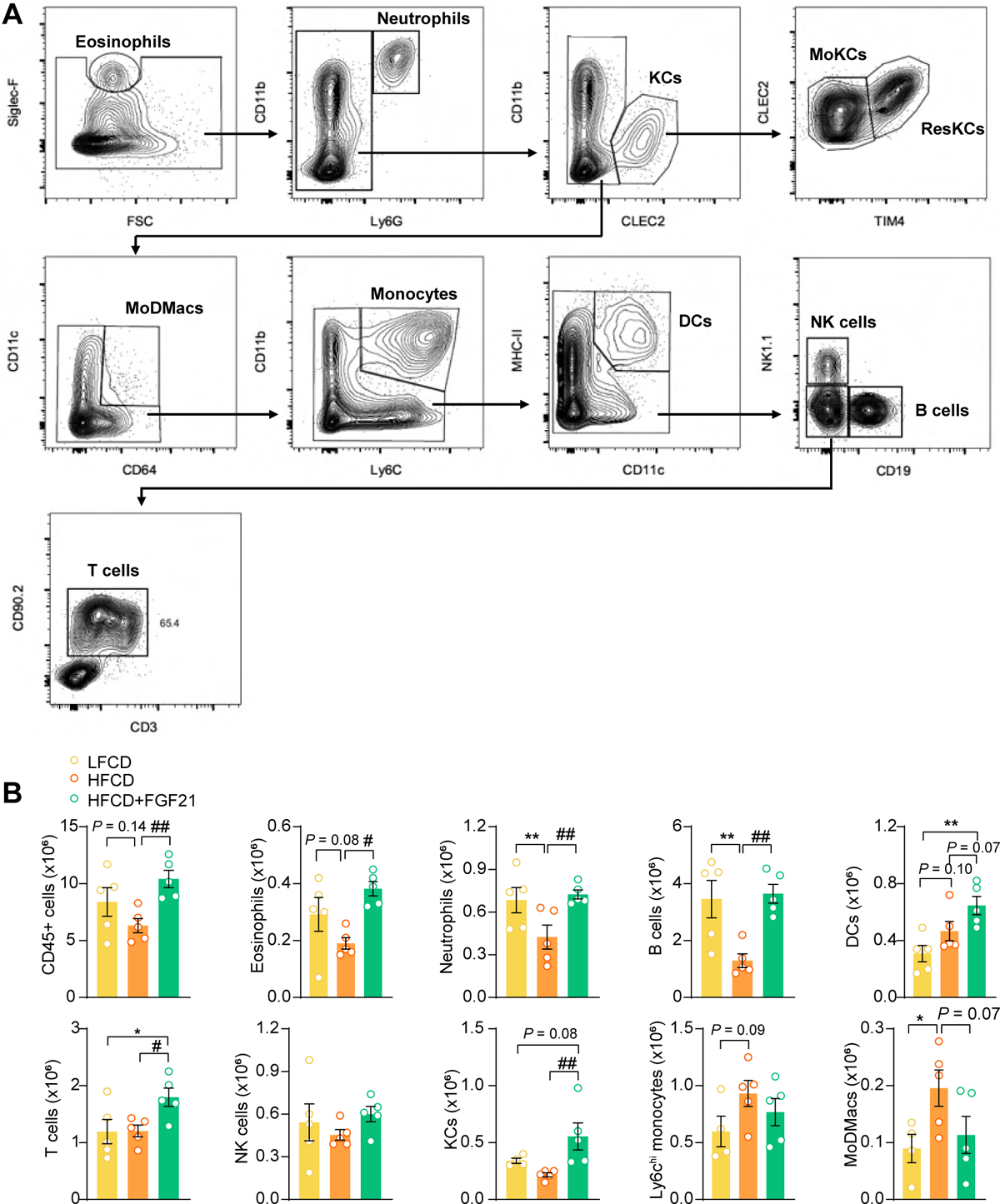
FGF21 modulates the hepatic immune cell pool. (**A**) Flow cytometry gating strategy. (**B**) After 23 weeks of treatment, CD45^+^ cells were isolated from the liver, and the number of CD45^+^ cells, eosinophils, neutrophils, B cells, dendritic cells (DCs), T cells, natural killer (NK) cells, total Kupffer cells (KCs), Ly6C^hi^ monocytes and monocyte-derived macrophages (MoDMacs) was assessed. Data are shown as mean ± SEM (n = 4-5 per group). Differences were assessed using one-way ANOVA followed by a Fisher’s LSD test. \**P* < 0.05, \*\**P* < 0.01, compared with the LFCD group. ^#^*P* < 0.05, ^##^*P* < 0.01, compared with the HFCD group.

**Figure 6-figure supplement 2.**
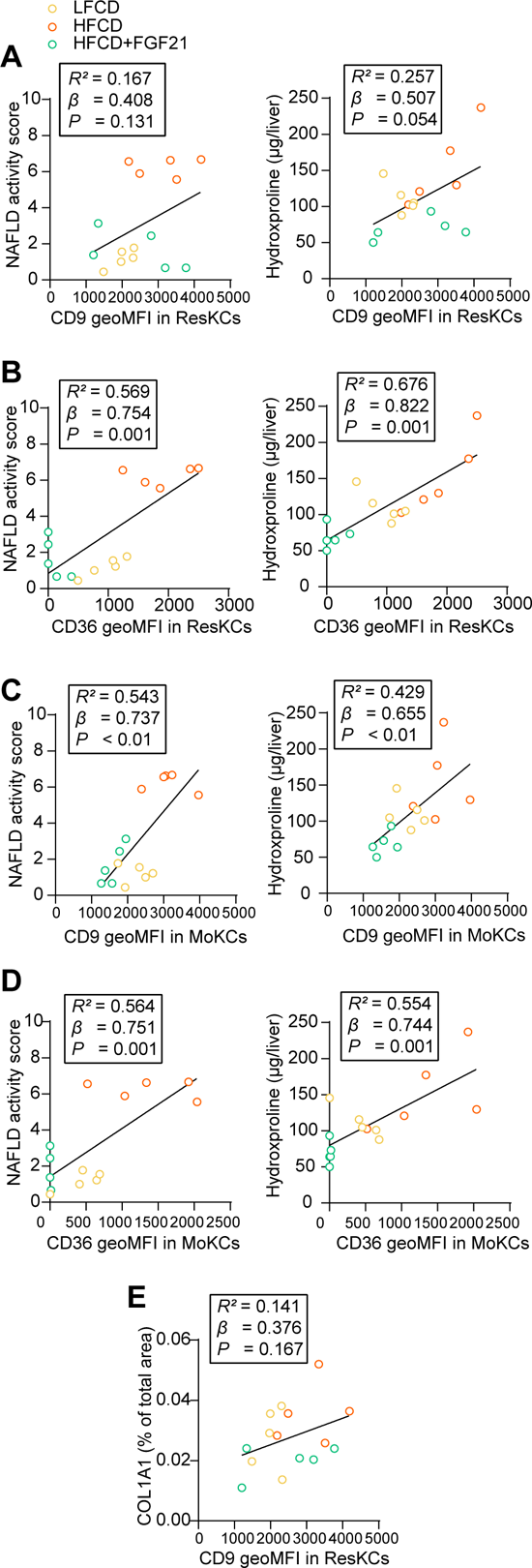
CD36^hi^ ResKCs as well as CD36^hi^/CD9^hi^ MoKCs positively correlate with NAFLD activity score and liver fibrosis. NAFLD activity scores and liver hydroxyproline levels were plotted against the expression of (**A**) CD9 and (**B**) CD36 in ResKCs as well as (**C**) CD9 and (**D**) CD36 in MoKCs. (**E**) Hepatic expression of collagen type 1α 1 (COL1A1) was plotted against the expression of CD9 in ResKCs. Linear regression analyses were performed. Data are represented as mean ± SEM (n = 5 per group).

## List of Supplementary Files

**Supplementary File 1:** Supporting Materials and Methods.

---

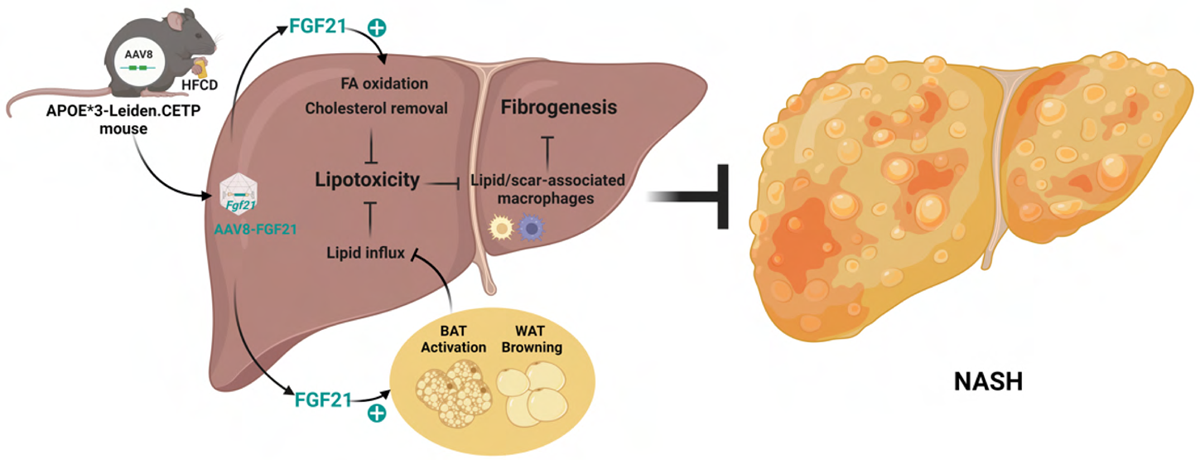

## Notes

Grant support: This work was supported by the Dutch Diabetes Research Foundation (2015.81.1808 to M.R.B.); Netherlands Organisation for Scientific Research-NWO (VENI grant 91617027 to Y.W.); Chinese Scholarship Council grants (CSC 201606010321 to E.Z.); the Novo Nordisk Foundation (NNF18OC0032394 to M.S.); and the Netherlands Cardiovascular Research Initiative: an initiative with support of the Dutch Heart Foundation (CVON-GENIUS-2 to P.C.N.R.).

